# Control of fluid flow by Adgrd1 is essential for mammalian oviductal embryo transport

**DOI:** 10.1101/2020.06.15.144444

**Authors:** Enrica Bianchi, Yi Sun, Alexandra Almansa-Ordonez, Michael Woods, David Goulding, Nadia Martinez-Martin, Gavin J. Wright

## Abstract

Dysfunction of oviductal embryo transport can lead to ectopic pregnancy which affects 1 to 2% of all conceptions in the United States and Europe, and is the most common cause of pregnancy-related death in the first trimester^1, 2^. Ectopic pregnancies almost always occur in the Fallopian tube, emphasizing the critical role of oviductal transport in human reproduction^3^. Oviductal transit is regulated and involves a valve-like “tubal-locking” phenomenon that temporarily arrests oocytes at the ampullary-isthmic junction (AIJ) where fertilization occurs^4^. Here, we show that female mice lacking the orphan adhesion G-protein coupled receptor *Adgrd1* are sterile because they are unable to unlock the restraining mechanism at the AIJ, inappropriately retaining embryos within the oviduct. *Adgrd1* is expressed on the oviductal epithelium and the post-ovulatory attenuation of tubal fluid production is dysregulated in *Adgrd1*-deficient mice. We identified Plxdc2 as an activating ligand for Adgrd1 displayed on the surface of cumulus cells. Our findings suggest that regulating oviductal luminal fluid production by Adgrd1 controls embryo transit, and provides important insights into the genetic regulation and molecular mechanisms involved embryo tubal transport.

## Main

Ectopic pregnancies occur when a fertilized egg implants and develops outside of the uterus, and in almost all cases, this occurs in the Fallopian tube resulting in a tubal pregnancy. The control of embryo movement through the oviduct is therefore thought to have an important role in this condition, but while generic risk factors that include previous IVF treatment and surgery have been identified, the underlying genetic causes and mechanisms are poorly characterised^3^. The oviduct promotes successful fertilization by storing and supporting the capacitation of sperm, and providing a conduit for eggs and early embryos to reach the uterus. The transit of eggs and embryos through the oviduct is thought to be due to the combined action of several different factors which include the beating of cilia in an abovarial direction, periodic contractions of the surrounding muscle, and regulated secretions from the oviductal epithelium. One general feature of mammalian oviductal transport is that ovulated cumulus-oocyte-complexes initially move rapidly through the infundibulum to the ampulla, and are then halted for many hours at the ampullary-isthmic junction (AIJ) before continuing their journey to the uterus^4–8^. The AIJ is the site of fertilization and the arrest of oocytes is likely to promote successful reproduction by providing a suitable environment for the gametes to meet and suppress polyspermy^9^. The pausing of tubal passage is a conserved feature of mammalian oviducts and has been described in several mammals including mouse^6, 7^, rabbits^8^, horses^10^, sheep, pig, guinea pig and cat^11^. None of the factors known to be involved in tubal transport provide a satisfactory explanation for this valve-like behaviour of the oviduct, and especially how it is “unlocked” to allow the developing embryos continued passage to the uterus.

To identify genes required for female fertility with unknown mechanisms of action, we interrogated the International Mouse Phenotyping Consortium database and identified a gene encoding an orphan member of the family of adhesion G-protein coupled receptors, *Adgrd1*^12^. We confirmed that female mice containing a targeted *Adgrd1* gene-trap allele (Supplementary Fig. 1) were sterile, irrespective of the male genotype (Fig. 1a). Female *Adgrd1*-mutant reproductive tissues were morphologically normal, ovulated the same number of eggs as wild-type littermates (Fig. 1b), and fertilization was unaffected both *in vivo* (Fig. 1c) and *in vitro* (Fig. 1d). No embryo implantation was observed in *Adgrd1*-deficient mothers (Fig. 1e), although *Adgrd1^-/-^* uteri could support embryo development (Fig. 1f, g) since transferred wild-type embryos developed normally until at least day 12.5 (Fig. 1g - inset). We consistently observed embryos denuded of cumulus cells that were ectopically located in the ampulla of mutant mothers (Fig. 1h, i), which had reached the expected morula stage by E2.5 (Fig. 1j). The remnants of unfertilized eggs such as empty zonae pellucidae and dead oocytes were also frequently observed in the ampullae of mutant mothers (Fig. 1h). Together, these data demonstrate that *Adgrd1^-/-^* females are sterile because embryos are inappropriately retained in the ampulla, perhaps because they are unable to reverse the “tubal lock” that retains eggs at the AIJ.

**Fig 1:**
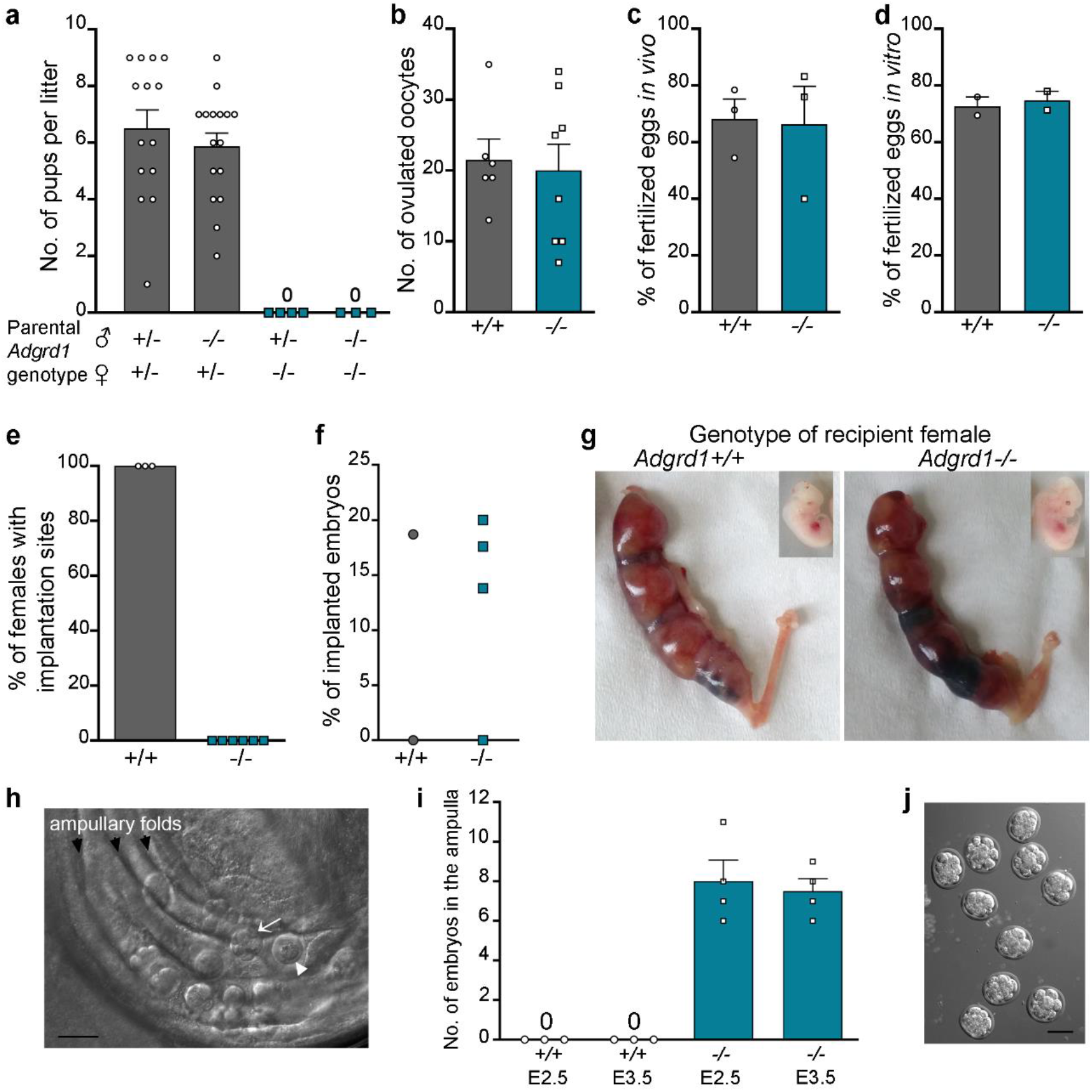
*Adgrd1^-/-^* female mice are infertile due to defective embryo transport. **a,** *Adgrd1^-/-^* females are sterile despite normal oocyte ovulation **b,** and fertilization **c** and **d**. **e,** No implantation sites were observed in *Adgrd1^-/-^* females. **f**, Mutant uteri support implantation of transferred embryos. **g**, Implantation of wild-type transferred embryos in *Adgrd1^-/-^* and control uteri at E12.5; embryos developed normally (inset). **h,** E2.5 embryos are ectopically located within the ampulla of an *Adgrd1^-/-^* oviduct. Arrow identifies morula-stage embryo, arrowhead identifies dead oocyte. Scale bar represents 100 μm. **i**, embryos are ectopically located in the ampulla of *Adgrd1^-/-^* mice at E2.5 and E3.5. **j**, Mutant *Adgrd1^-/-^* embryos recovered from the ampulla at E2.5 have reached the morula stage. Scale bar represents 80 μm. Bars in **a**, **b**, **c**, **d**, and **i** represent mean ± s.e.m.

To understand how loss of Adgrd1 regulates embryo passage through the AIJ, we first showed that *Adgrd1* was highly transcribed in the isthmus (Fig. 2a). The protein was localised at the surface of both secretory and ciliated oviductal epithelial cells (Fig. 2b and Supplementary Fig. 2). Consistent with similar experiments in other mammals^8, 13^, tissue sections of wild-type and *Adgrd1^-/-^* oviducts throughout the estrus cycle failed to reveal a constriction or occlusion at the AIJ that could explain embryo retention, and this was consistent with the ease of flushing trapped embryos from the ampulla of *Adgrd1^-/-^* oviducts through the isthmus. Embryo transport through the oviduct is thought to involve the concerted effects of oviductal fluid secretions, muscle contractions, and the action of cilia which beat in an abovarial direction towards the uterus, although their relative contributions have not been determined. We did not find any evidence that the morphology or function of muscle contractions were disrupted in *Adgrd1^-/-^* oviducts (Supplementary Fig. 3a. Supplementary Movies 1 and 2). Video microscopy showed that beads placed into explants of Adgrd1-mutant oviducts were not rigidly held in position, but rather were moved back and forth within the lumen of the oviduct in pendulum-like movements due to the contractions of the surrounding muscle (Supplementary Movie 1). These observations were consistent with the view that muscle contractions mix rather than vectorially transport the contents of the tube^11^. Similarly, we did not find any overt difference in either the number, function or, by electron microscopy, ultrastructure in cilia of *Adgrd1^-/-^* oviducts compared to wild-type controls (Supplementary Fig. 3b, c, d and Supplementary Video 2). This finding was consistent with the transport of the cumulus complexes from the infundibulum to the ampulla which was normal in *Adgrd1^-/-^* mice, and thought to be largely driven by ciliary action.

**Fig 2:**
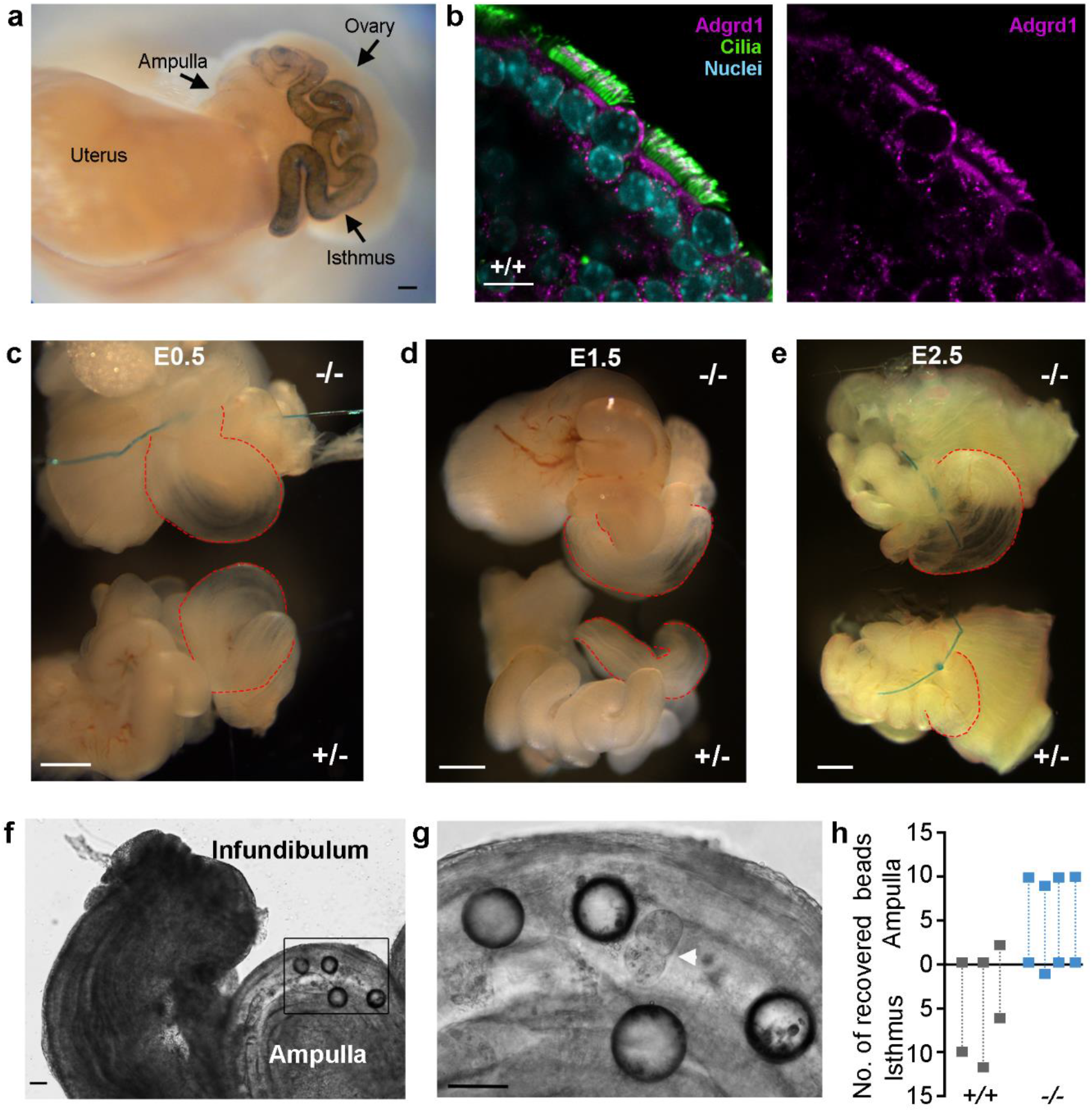
*Adgrd1* is expressed on the oviductal epithelium and regulates post-ovulatory attenuation of oviductal fluid production. **a**, X-gal staining of *Adgrd1^+/-^* female reproductive tract shows strong *Adgrd1* promoter activity in the isthmus. Scale bar represents 200μm. **b**, Adgrd1 is localized on the apical membrane of ciliated and non-ciliated cells in the ampullary oviductal epithelium. Scale bar represents 10μm. **c**, Accumulation of fluid is comparable in ligated control (+/-) and mutant (-/-) oviductal explants *in vitro* at E0.5; ampullary distension is marked with red lines. **d**, No reduction in fluid accumulation in mutant *Adgrd1^-/-^* oviducts at E1.5 by contrast to control. **e**, Oviductal ligation at E2.5 *in vivo:* more fluid has accumulated in the ampulla of the mutant compared to control; a representative example is shown. Images in **c**, **d** and **e** are representative of a least three independent experiments and scale bar represents 500μm. **f**, Glass beads (~100μm diameter) were inappropriately retained in the ampulla of *Adgrd1^-/-^* oviducts. Image taken 24 hours after bead transplantation. **g**, Magnification of the boxed area in **f,** showing glass beads and retained embryo (arrowhead). Scale bars represent 80 μm. **h,** Distribution of 10 to 12 transplanted glass beads in *Adgrd1^-/-^* (*n* = 4) and control (*n* = 3) oviducts at E1.5.

Oviductal fluid influences embryo transport and varies throughout the estrus cycle, peaking at ovulation and reducing in volume prior to the next cycle^14–17^. Due to the small size and flexuous morphology of the mouse oviduct, ligation is a practical method of evaluating fluid production^18^, and so heterozygous *Adgrd1^+/-^* oviductal explants were ligated at the infundibulum *in vitro* at E0.5 and E1.5. Four hours after ligation of E0.5 oviducts, we observed a striking distention of the ampulla due to the accumulated fluid (Fig. 2c). This demonstrated that fluid exits the oviduct at the ovarial end creating a flow that opposes the movement of embryos - agreeing with observations in mice^18^, and larger mammals^13, 15, 19, 20^. Consistent with the reduction in fluid production after ovulation, ligating heterozygous oviducts in the post-ovulatory period (E1.5) resulted in a much reduced accumulation of fluid (Fig. 2d). *Adgrd1^-/-^* mutant oviducts ligated at E0.5 exhibited the same ampullary distention (Fig. 2c); however, in contrast to heterozygous littermates, did not show the same reduction at E1.5 demonstrating that the post-ovulatory attenuation of fluid production in mutant oviducts is dysregulated (Fig. 2d). This was confirmed by performing the ligations *in vivo* at E2.5, which resulted in a more conspicuous distension of the ampulla due to the accumulated fluid in the mutant compared to heterozygous controls (Fig. 2e and Supplementary Figs. 4a and b). Dysregulated fluid production was specifically observed in the isthmus by ligating oviducts within the isthmus itself (Supplementary Fig. 4c). Together, these results show that the postovulatory cessation of oviductal fluid production is misregulated in the isthmus of Adgrd1-deficient mice which could impede embryo transport and cause infertility. We therefore reasoned that if embryo retention was due to dysregulated oviductal fluid production, the transport of particles other than embryos would be affected. Consistent with this, we found that appropriately-sized glass beads were similarly retained within the ampulla of *Adgrd1^-/-^* but not control oviducts (Fig. 2f, g and h).

Adgrd1 encodes a cell surface receptor belonging to the adhesion G-protein coupled receptor family, a subset of over 30 proteins that typically contain a large N-terminal ectodomain, most of which have no identified ligand^21^. Adgrd1 contains a pentraxin domain in its extracellular region and is known to initiate intracellular signalling by stimulatory G proteins leading to increases in cAMP levels by activating adenylate cyclase^22^. One mechanism to trigger G-protein signalling used by this family of receptors is through the ligand-dependent relief of an auto-inhibitory ectodomain^21, 23, 24^. To identify a ligand for ADGRD1, we first expressed the entire ectodomain as a multimeric binding probe to increase binding avidity and thereby circumvent the often weak binding affinities of extracellular receptor-ligand interactions^25^. The highly avid ADGRD1 binding probe was then systematically tested in an unbiased manner for binding to a panel of 1,132 unique human receptor ectodomains^26^, and the Plexin Domain-Containing Protein 2 (PLXDC2) was identified as a candidate ligand (Supplementary Fig. 5a). To investigate this further, we used an assay designed to detect extracellular receptor-ligand interactions called AVEXIS^27^ and demonstrated that ADGRD1 and PLXDC2 interacted in both bait-prey orientations (Fig. 3a). To further validate the interaction, we expressed the entire ectodomains of the mouse orthologues of both Adgrd1 and Plxdc2, and showed that they could interact (Fig. 3a). Cells transfected with a plasmid encoding Plxdc2 gained the ability to bind the highly avid Adgrd1 binding probe, and this binding was specifically blocked by preincubating the transfected cells with an anti-Plxdc2 antibody (Fig. 3b). We next mapped the interaction interface to the pentraxin domain of Adgrd1 and the PSI domain of Plxdc2 by creating a series of truncated ectodomains that encompassed known domains of both proteins and using the AVEXIS assay to quantify binding (Fig. 3c, d and Supplementary Fig. 5b). To understand how the Adgrd1 receptor could be activated in the oviduct, we characterised the tissue expression patterns of Plxdc2. Within the female reproductive system, *Plxdc2* was most highly transcribed in the ovary and cumulus-oocyte complexes (COCs) compared to the uterus and oviduct and other tissues such as brain, liver and spleen (Supplementary Fig. 5c). We conformed the expression of Plxdc2 protein in the ovary and COCs by Western blotting (Fig. 3e), and showed using immunocytochemistry that Plxdc2 was highly expressed on the surface of cumulus cells (Fig. 3f). We demonstrated that Plxdc2 is an activating ligand for Adgrd1 using an established GPCR activation assay which measures the increase in cellular cAMP levels^28^. *Adgrd1*-transfected HEK293 cells showed an expected increase in cAMP levels^29^ which was augmented by Plxdc2 (Fig. 3g). Previous work in larger mammals including humans showed that addition of cell permeable dibutyryl cAMP and agents that increase cAMP levels such as forskolin, theophylline and cholera toxin all resulted in a decrease or abolition of oviductal secretion^14, 30, 31^.

**Fig. 3:**
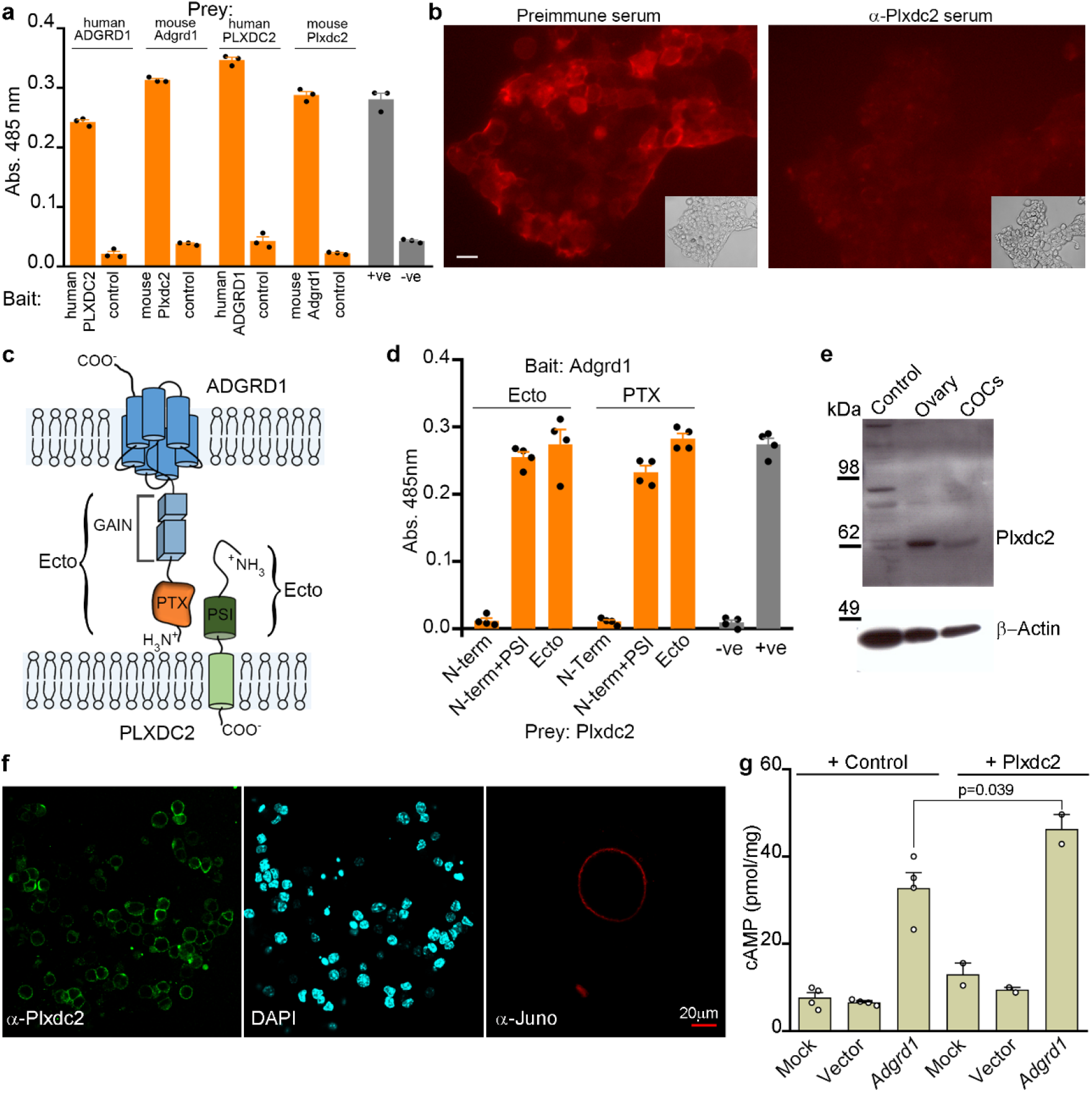
Plxdc2 is an activating ligand for Adgrd1 and expressed on cumulus cells. **a,** Direct interactions between human and mouse ADGRD1 and PLXDC2 ectodomains in both bait-prey orientations using the AVEXIS assay. **b,** Adgrd1 ectodomain probe bound *Plxdc2*-transfected cells (left panel) and was blocked by preincubation with Plxdc2 antiserum (right). Scale bar represents 10 μm. **c**, Schematic of ADGRD1 and PLXDC2 domain organisation. **d**, Domain truncations demonstrate that the pentraxin (PTX) domain of Adgrd1 and PSI domain of Plxdc2 are sufficient for binding. **e**, Western blotting and immunofluorescence **f**, demonstrates that Plxdc2 is expressed on the plasma membrane of cumulus cells. Anti-Juno shows the localization of the oocyte. **g**, The overexpression of *Adgrd1* induces a significant increase of intracellular cAMP compared to non-transfected cells (mock) and to cells transfected with a plasmid encoding GFP (vector). The Plxdc2 ectodomain induces a significant increase of cAMP levels in *Adgrd1*-expressing cells only. A representative of four independent experiments is shown. Bars in **a**, **d**, and **g** represent mean ± s.e.m.

Our results show that Adgrd1-mediated control of oviductal fluid production provides a mechanism to explain the valve-like behaviour of the mammalian oviduct. We present a model (Fig. 4) where ovulated COCs are propelled towards the uterus by ciliary action but are halted at the AIJ due to the opposing force of oviductal fluid flowing towards the ovary caused by the narrowing of the oviduct at the isthmus. Adgrd1 is locally activated in the oviductal epithelium by Plxdc2 displayed on the surface of cumulus cells that are progressively released as the jelly-like hyaluronic acid matrix surrounding the cumulus mass gradually disintegrates. The triggering of Adgrd1 decreases oviductal fluid production and consequently the flow that opposes the constitutive abovarial ciliary action to permit continued passage of embryos through the isthmus. This suggests that cumulus cells not only support oocyte development but also communicate with the oviduct to regulate tubal transit. Our demonstration that Adgrd1 is essential for embryo transport provide new avenues for the diagnosis and treatment of ectopic pregnancy which is a major risk for human reproductive health.

**Fig. 4:**
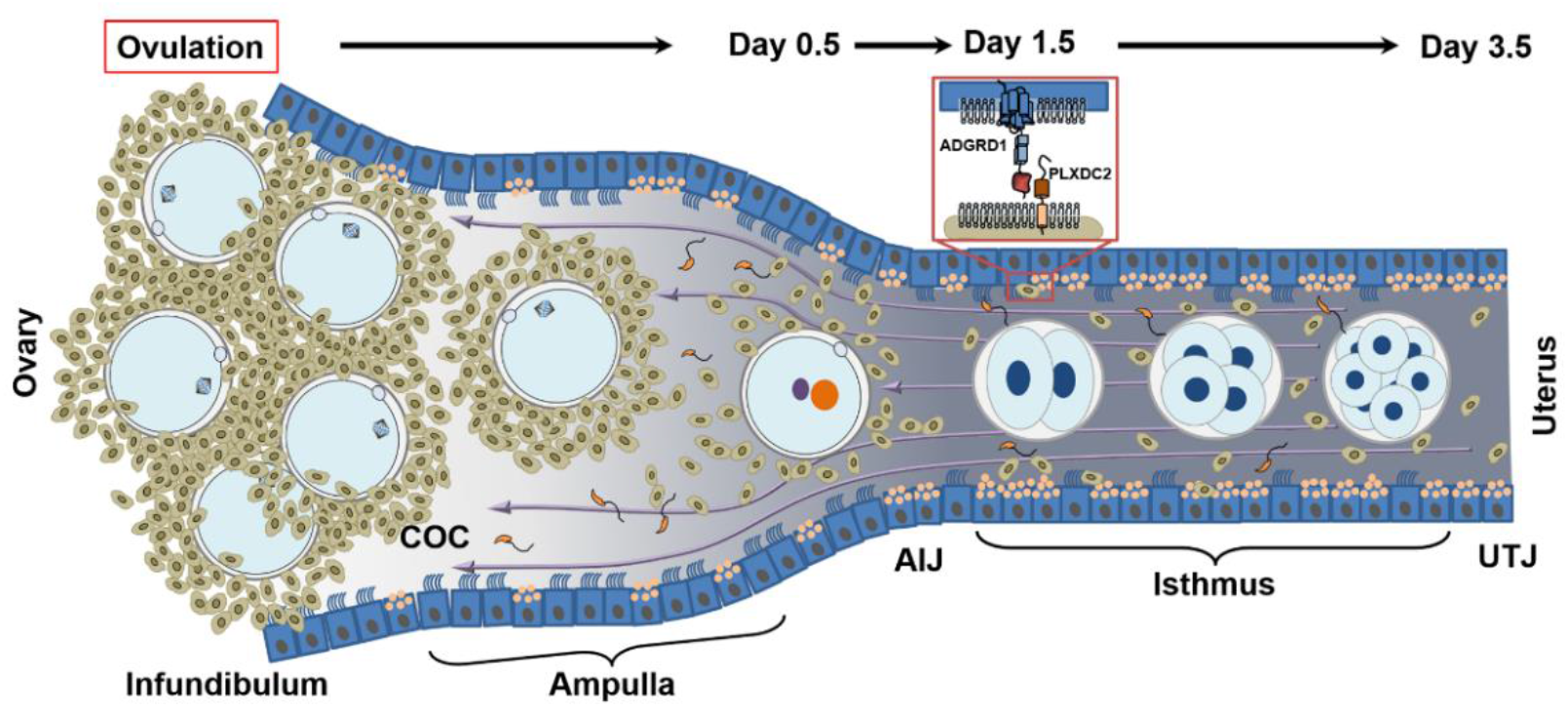
Model describing Adgrd1-mediated control of fluid flow underlies the valve-like behaviour of the mammalian oviduct. Cumulus-oocyte-complexes are ovulated into the infundibulum and rapidly transported to the AIJ by the action of the cilia lining the oviductal epithelium beating towards the uterus. At the AIJ, the COCs are halted due to the balance of abovarial ciliary action and the force of adovarial oviductal fluid flow (purple arrows). In wild-type oviducts, the gradual release of cumulus cells as the hyaluronic acid matrix which surrounds them slowly disintegrates releases the Plxdc2 ligand and triggers a reduction of oviductal fluid production by activating Adgrd1 on the oviductal epithelium to licence embryo release. COC = cumulus-oocyte-complex; AIJ = ampullary-isthmic junction; UTJ = uterotubal junction.

## Acknowledgments

We thank the Sanger Mouse Genetics Programme for generating the *Adgrd1^-/-^* mice and for the initial phenotyping; the staff at the Sanger Research Support Facility for the provision and help with the management of the mice. This work was funded by the Medical Research Council (grant MR/M012468/1) to EB and GJW and Wellcome Trust (grant 206194) to GJW.

## Author contributions

EB performed all experiments unless stated. Mouse surgery was performed by MW; Receptor screening was performed by YS and NM-M; domain mapping experiments were performed by AA-O; electron scanning and transmission microscopy was performed by DAG. The manuscript was written by EB and GJW.

## Materials and Methods

### Generation, breeding and fertility phenotyping of *Adgrd1*-deficient mice

All animal experiments were performed under UK Home Office governmental regulations and European directive 2010/63/EU. Research was approved by the Sanger Institute Animal Welfare and Ethical Review Board. *Adgrd1^-/-^* mice were obtained from the Knockout Mouse Project resource (IKMC Project: 22527) and contain a lacZ-tagged allele targeted to the *Adgrd1* genomic locus located on chromosome 5, *Adgrd1^tm1b(EUCOMM/Wtsi)^*. Mice were generated by injecting blastocysts with the targeted mouse embryonic stem cells which were transferred to pseudopregnant females to generate chimeras. Germline transmission of the targeted allele was confirmed by PCR after mating of chimeric males with C57BL/6NTac females. To obtain the reporter-tagged deletion allele, females heterozygous for the ‘knockout-first’ allele *Adgrd1^tm1a(EUCOMM/Wtsi)^* were crossed to hemizygous males ubiquitously expressing the Cre enzyme^32^. Mice with the recombined allele were identified using PCR and diagnostic primers using genomic DNA extracted from ear biopsies (4403319 DNA Extract All Reagents Kit, Life Technologies) as the template for short range PCR using Platinum^®^ Taq DNA Polymerase (10966034 Invitrogen). Wild-type and mutant alleles were amplified using the same sense primer: 5’-ACTTTGTGGGTGGTGTCCG-3’ and two specific anti-sense primers, 5’-CGTTCATGCAAGCCATCAC-3’ for the wild-type and 5’-TCGTGGTATCGTTATGCGCC-3’ for the mutant recombined allele. The mouse colony was maintained by crossing heterozygous males and females. Male and female fertility was quantified by pairing homozygous and heterozygous *Adgrd1* transgenic adult mice with homozygous and heterozygous animals of proven fertility and the number of resulting pups was monitored continuously for three months. The number of ovulated oocytes was counted after induction of ovulation with 5 IU of pregnant mare serum gonadotropin (PMSG) followed by 5 IU of human chorionic gonadotropin (hCG) 48 hours later. *In vivo* fertilization was assessed by scoring the number of zygotes and the number of non-fertilized oocytes present in the ampulla at embryonic day 0.5. Eggs and embryos were fixed in 4% formalin (28906 Thermo Scientific Pierce) and stained with DAPI (62248 Thermo Fisher). *In vitro* fertilization was performed essentially as described^33^. Briefly, sperm were collected from the cauda epididymis of adult male mice, capacitated for 1 hour in HTF medium at 37 °C and added to cumulus-enclosed oocytes that had been collected 13 hours after hCG treatment. Groups of 20 to 30 eggs were inseminated in 100 μL drops of HTF medium containing 1 x 10^5^ sperm; four hours later, eggs were washed and cultured in KSOM (MR-121D EmbryoMax, Millipore) for the following four days. Formation of pronuclei was scored and embryo development recorded daily. Mutant and age-matched control females were mated with proven males and the day of vaginal plug formation was counted as E0.5. The female reproductive tracts were visually inspected for the presence of implantation sites at E5.5, E6.5 and E8.5. The embryo position in the oviducts was determined using the Zeiss Axioplan 2 microscope and embryos were counted after flushing the oviducts with M2 medium (Sigma).

Non-surgical embryo transfer (NSET) was performed as described^34^. Wild-type 2-cell embryos were cultured in KSOM for 48 hours and blastocysts were transferred to pseudopregnant *Adgrd1^+/+^* and *Adgrd1^-/-^* recipient females at E2.5 using the NSET device (Paratechs, catalogue #60010). 14 and 16 blastocysts were transferred to two wild-type females, and 14, 17, 29 and 30 blastocysts were transferred to four *Adgrd1*-deficient females. The number of implanted embryos was counted ten days after the transfer.

### Time-lapse video microscopy

Oviducts were carefully dissected from 8-week-old females at the required stage of the estrus cycle. Entire oviducts were separated from the ovary by opening the ovarian bursa, and scission of the ovarian ligament; isolation from the rest of the reproductive tract was achieved by trimming the broad ligament whilst maintaining the utero-tubal junction. When cutting the mesosalpinx, care was taken to avoid pulling and damaging the tubes. Spontaneous rhythmic contractions of oviducts were documented by video recording immediately after dissection using a Zeiss Axioplan 2 microscope equipped with a Hamamatsu 1394 ORCA-ERA digital camera and the Velocity imaging software. Fifty seconds of real time were acquired at a frame frequency of 1 Hz. To record ciliary beating, the ampullary region of the oviducts were opened longitudinally, placed in PBS on a microscope glass and microparticles (Sigma 74964) were gently added to the tissue. Movies were acquired at 8.5 frames per second.

### Wholemount LacZ staining, immunofluorescence and immunohistochemistry

To determine the expression of the lacZ reporter gene in the targeted allele at the *Adgrd1* locus, dissected tissues were fixed in 10% Neutral Buffered Formalin (NBF, Cellpath) overnight, washed with PBS, stained with a LacZ staining solution (2 mM MgCl2, 0.02% IGEPAL CA-630, 5 mM potassium ferrocyanide, 5 mM potassium ferricyanide, 0.01% deoxycholic acid, 0.1% X-Gal (5-bromo-4-chloro-3-indolyl β-D-galactopyranoside) in dimethylformamide) at 4 °C, post-fixed in 4% formalin and stored in 70% glycerol.

Polyclonal antiserum was raised by immunizing rabbits with the entire ectodomains of mouse ADGRD1 or mouse PLXDC2 expressed as soluble recombinant 6-his-tagged proteins in HEK293-6E cells and purified using Ni^2+^-NTA chromatography, essentially as described^35^. Immunizations were carried out by Cambridge Research Biochemicals in accordance with UK Home Office regulations and the Sanger Institute animal welfare ethical review board.

To prepare wholemount tissues for immunofluorescence, they were fixed in 4% formalin for 30 minutes at room temperature, incubated with the primary antibody for 1 hour, and washed extensively with PBS and finally stained with the secondary antibody conjugated with the appropriate fluorochrome (Alexa Fluor 488 goat anti-mouse supplied by Thermo Fisher, and Alexa Fluor 647 goat anti-rabbit supplied by Jackson Immunoresearch). Where tissues were sectioned, tissue was fixed in 4% formalin for 60 minutes, rinsed extensively in PBS and soaked in a sucrose solution overnight before embedding in OCT and 8 micron sections cut with a Leica CM1950 cryostat. Sections were treated with 1% SDS in PBS for 5 minutes, blocked before adding primary antibody overnight at 4 °C. Sections were washed and incubated with a goat anti-rabbit – Alexa488 secondary antibody for 1 hour at room temperature. Sections were then washed and mounted in slowFade Gold mounting solution with DAPI (Thermo).

Binding of recombinant proteins to cells was quantified by incubating transfected cells with highly avid ectodomains expressed as FLAG-tagged pentameric preys in cell culture medium at 37 °C before fixation in a phosphate-buffered 4% formalin solution. Staining of muscle was performed by first permeabilizing fixed oviducts with 0.2% Triton X-100 and incubating with Texas-Red-phalloidin conjugate for 30 minutes. Optical sections of whole oviductal tissue were acquired with a Leica SP5 laser confocal microscope (z-step size = 2.52μm). The 3D projection of the whole z-stack was generated with the LAS AF software. The antibodies and probes used in this study were: polyclonal rabbit-anti-mouse Adgrd1 (this study); polyclonal rabbit-anti-mouse Plxdc2 (this study); rat monoclonal anti-mouse Juno (clone TH6, Biolegend); mouse monoclonal anti-acetylated tubulin (clone 6-11-B-1, Sigma Aldrich); mouse monoclonal anti-Flag-Cy3 conjugate (clone M2, Sigma Aldrich); polyclonal rabbit anti-beta-actin (Abcam, ab8227); Texas-red conjugated phalloidin (Thermo Fisher, T7471); -; anti-ADGRD1 (rabbit polyclonal, HPA042395, Atlas antibodies).

### Transmission and scanning electron microscopy

Mouse oviducts were fixed at room temperature in a 2% formalin / 2.5% glutaraldehyde mixture buffered in 0.1 M sodium cacodylate at pH 7.4 for one hour, rinsed and fixed in 1% osmium tetroxide for another hour followed by 1% tannic acid and 1% sodium sulphite for 30 minutes respectively. Samples were then dehydrated in an ethanol series, staining en bloc with uranyl acetate at the 30% stage before embedding (Epoxy embedding medium kit – Sigma). 50 nm ultrathin sections were cut on a Leica UC6 ultramicrotome and imaged on an FEI 120kV Spirit Biotwin TEM with a F4.15 Tietz camera.

For scanning electron microscopy, mouse oviducts were fixed as for TEM (replacing tannic acid with osmium-thiocarbohydrazide), dehydrated in an ethanol series, critical point dried in a Leica CPD300, mounted and sputter-coated with 2 nm of platinum using a Leica ACE600 evaporator. Samples were imaged on a Hitachi SU8030 SEM.

### Oviduct ligation, surgical procedures and transfer of beads

Oviducts from mice at the required stage in the estrus cycle were dissected under a Nikon SMZ800 stereo microscope. Once the ovary was separated from the oviduct and removed, a surgeon’s knot was tied around the oviduct with polypropylene suture (Ethicon Prolene 10-0, W2794). The ligated oviducts were maintained in culture at 37 °C in 500 μL of pre-warmed KSOM media, fixed in NBF, and imaged with a Leica M205FA stereomicroscope.

Glass beads with a diameter of ~100 μm (Sigma G4649) for transfer into oviducts were sterilised by autoclaving and washed in M2 medium. Female recipient mice whose mating had been timed were prepared for surgery using gaseous anaesthesia under aseptic conditions, shaved, and given analgesia. Using a stereomicroscope (Leica MZ75), a small longitudinal skin incision was made at the midline, level with the last rib. The ovary was located through the muscle wall and micro scissors were used to make a small incision, followed by blunt dissection and exteriorisation of the ovary and oviduct by holding the associated fat pad, anchored using a serrefine clamp. After removing the bursa covering the ovary and oviduct, glass beads were taken up using a mouth pipette into a 230 mm Pasteur glass pipette with the aid of a stereomicroscope (Leica MZ95) and transferred into the oviducts of recipient mice at E0.5. The location of the beads in the ampulla and isthmus were counted at E1.5. *In vivo* ligation of oviducts was performed using the same surgical procedure with polypropylene suture which was slid under the oviduct ~1mm from the infundibulum and fine forceps used to tie a surgeon’s knot. When assessing fluid accumulation in the isthmus, additional ligatures were tied at the ampullary-isthmic junction and the isthmus. The ovary and oviduct were carefully replaced within the body cavity using blunt forceps, and the skin incision closed using a single wound clip.

### Preparation of the protein library for large-scale receptor screening

A library of single-transmembrane-spanning (STM) human proteins was compiled using computational prediction algorithms and experimental evidence, as described^26^. The ectodomain of each protein was synthesized and cloned into the pRK5 vector (Genentech) in frame with a C-terminal Fc (hIgG1) tag. The resulting library consists of 1,364 human proteins, including 1,132 unique receptors and a number of replicate constructs as assay controls. Expression of the human STM protein library was performed as described^26^. In brief, conditioned media enriched in individual receptors was prepared in Expi293F cells (Thermo Fisher), transiently transfected with receptors expressed as ectodomains fused to a human Fc (IgG1). Expi293F cells were cultured in Expi293 Expression Media (Cell Technologies) in flasks at 37 °C and 150 rpm agitation in a humidified incubator. Human cells were chosen for expression of the collection of STM receptors to maximize protein quality and incorporation of relevant post-translational modifications. One mL cell transfections were performed using an automated system consisting of a TECAN liquid handling system and a MultiDrop reagent dispenser. Transfections were processed in batches of 96 clones using a Biomek FX liquid handling robot and conditioned media was harvested 7 days post-transfection. Subsequently, the receptor Fc-tagged ectodomains present in the conditioned media were captured on protein A-coated 384-plates (Thermo Scientific), and stored at 4 °C until use. For the binding screen, the ectodomain of ADGRD1, truncated at S600, was expressed in HEK293-6E cells as a pentameric probe fused to a beta-lactamase enzyme^27^. A Thr to Gly mutation at position 574 in the GPCR proteolytic site (GPS) of human ADGRD1 was introduced to prevent the autoproteolysis^36^.

### Identification of binding extracellular ligands for ADGRD1

The Cell Surface Receptor Interaction screening was based on the AVEXIS method^27^, which was further implemented for automated high throughput screening in 384 well plate format, as recently described^26^. In brief, screens were performed using an integrated robotic system consisting of automated liquid handling devices for high throughput analysis of receptor-ligand interactions. On the day of the assays, the protein A plates coated with the receptor library were washed with PBS containing Ca^2+^ and Mg^2+^, followed by incubation with the beta-lactamase-tagged pentameric ADGRD1 probe for 1 hour at room temperature. Plates were then washed to remove free protein prior to addition of nitrocefin (Calbiochem) for 1 hour at room temperature. Hydrolysis of the beta-lactamase substrate nitrocefin was quantified by measuring absorbance at 485 nm using an integrated TECAN plate reader. The raw enzymatic absorbance values were analysed to calculate Z-scores across all proteins in the library. Comparisons to previous screens of this type identified immobilised ligands which repeatedly gave positive binding signals irrespective of the binding probes used, and included known lectins (e.g. SIGLEC family members and MRC1) which are likely to directly interact with common glycans presented on the binding probe.

### Recombinant protein production and protein interaction screening

The extracellular domains of ADGRD1 and PLXDC2 were expressed as soluble secreted proteins by transient transfection in HEK293-6E cells^27^. NCBI reference sequences for the expressed proteins were: mouse Adgrd1: NP_001074811; human ADGRD1: NP_001317426; mouse Plxdc2: NP_080438; human PLXDC2: NP_116201. The region encoding the entire predicted ectodomain was chemically synthesized by gene synthesis (GeneArt, Life Technologies) flanked by unique NotI, AscI restriction sites to facilitate cloning into mammalian expression plasmids to produce enzymatically biotinylatable monomeric ‘baits’ or pentameric FLAG-tagged ‘preys’; both bait and preys contained a rat Cd4d3+4 epitope tag as described. Biotinylated proteins were produced by co-transfecting a plasmid encoding a secreted version of the BirA enzyme^27^. To produce recombinant proteins corresponding to specific regions of both ADGRD1 and PLXDC2 ectodomains, primers were designed that would amplify the appropriate region from the plasmid encoding the entire ectodomain by PCR and the products cloned into both bait and prey expression plasmids using unique NotI and AscI sites^35^. Bait proteins were normalized by ELISA using a monoclonal antibody (Ox68) recognizing the rat Cd4d3+4 tag. Prey proteins were normalized to their β-lactamase activity to levels suitable for the AVEXIS assay as described^27^. Biotinylated baits that had been either purified or dialysed against PBS to remove excess free D-biotin were immobilised in streptavidin-coated 96-well microtitre plates (NUNC). Preys were incubated for one hour, washed three times in PBS/0.1% Tween-20 and once in PBS. Finally, 60 μl of 125 μg/ml nitrocefin was added and the absorbance measured at 485 nm on a Pherastar plus (BMG laboratories). A biotinylated protein consisting of just the rat Cd4d3+4 tag alone was used as a negative control bait and a biotinylated Ox68 monoclonal antibody (anti-prey) was used as a positive control. The rat Cd200-Cd200R interaction was used as a positive control.

### Western Blot

To perform Western blotting, proteins from mouse ovary, ovulated COCs, and HEK 293-T cells as negative control, were extracted with RIPA buffer and quantified with Bradford assay (Thermo) following manufacturer’s instructions. Briefly, normalized protein amounts were resolved under non-reducing conditions by SDS–PAGE and blotted to Hybond-P PVDF membrane (GE Healthcare) for 1 hour at 30 V. After blocking for 1 h with 2% BSA, the membrane was incubated for 1 hour with PLXDC2 antiserum BSA or with anti beta-Actin diluted 1:5000 in PBST/2% BSA, washed three times and incubated with a horseradish peroxidase (HRP)-conjugated anti-rabbit antibody. Proteins were detected using SuperSignalWest Pico Chemiluminescent substrate (Thermo Scientific) and developed on photographic film.

### RT-PCR

Total RNA was extracted from mouse tissues using Trizol reagent (Invitrogen) as per manufacturer’s instructions, resuspended in water, and quantified with a Nanodrop 1000 spectrophotometer (Thermo Scientific). SuperScript III reverse transcriptase (Invitrogen) was used to produce cDNA from 1 μg of RNA and subsequent amplification was obtained with the KOD hot start DNA polymerase (Novagen). *Plxdc2* was amplified with the following primers: 5’-ATCCAGGTGAAAGTCGGGTTG-3’ and 5’-GGGAGGTCGTGGTAGTTTGA-3’ annealing to two different exons and generating a 372bp long amplicon. For *beta-actin* the primers were: 5’-ATATCGCTGCGCTGGTCGTC-3’ and 5’-AGGATGGCGTGAGGGAGAGC-3’, and the amplicon size 516 bp. PCR products were resolved on a 1.5% agarose gel and imaged with an Azure c-600 gel documentation system.

### cAMP ELISA

Adherent HEK293-T cells were transiently transfected with either a plasmid encoding mouse *Adgrd1*, or a negative control encoding membrane-tethered EGFP (vector), or were treated with the transfection reagent Lipofectamine 2000 alone (mock). Twenty-four hours after transfection, cells were seeded on a streptavidin-coated microtitre plate that had been pre-incubated with the mouse PLXDC2 or control rat Cd200 biotinylated ectodomains. After three hours at 37 °C, the levels of cyclic AMP were determined in duplicate using a cAMP ELISA kit (ADI-900-066, Enzo Life Science) according to the manufacturer’s protocol, and normalized to the total protein concentration determined using the Bradford assay (ThermoFisher Scientific). One-way ANOVA showed a significant variation among conditions (F_5, 12_ = 43, p< 0.0001) and a post-hoc Bonferroni test indicated that intracellular cAMP differed significantly in *Adgrd1*-transfected cells treated with Plxdc2 compared to *Adgrd1*-transfected cells treated with the control protein CD200 (p=0.0391).

## References

1. Barnhart, K.T. Clinical practice. Ectopic pregnancy. N Engl J Med 361, 379–387 (2009).

2. Shaw, J.L., Dey, S.K., Critchley, H.O. & Horne, A.W. Current knowledge of the aetiology of human tubal ectopic pregnancy. Hum Reprod Update 16, 432–444 (2010).

3. Corpa, J.M. Ectopic pregnancy in animals and humans. Reproduction 131, 631–640 (2006).

4. Knobil, E. & Neill, J.D. The Physiology of reproduction, Edn. 2nd. (Raven Press, New York; 1994).

5. Pauerstein, C.J. & Eddy, C.A. The role of the oviduct in reproduction; our knowledge and our ignorance. J Reprod Fertil 55, 223–229 (1979).

6. Croxatto, H.B. & Ortiz, M.E. Egg transport in the fallopian tube. Gynecol Invest 6, 215–225 (1975).

7. Humphrey, K.W. Observations on transport of ova in the oviduct of the mouse. J Endocrinol 40, 267–273 (1968).

8. Greenwald, G.S. A study of the transport of ova through the rabbit oviduct. Fertil Steril 12, 80–95 (1961).

9. Coy, P., Garcia-Vazquez, F.A., Visconti, P.E. & Aviles, M. Roles of the oviduct in mammalian fertilization. Reproduction 144, 649–660 (2012).

10. Weber, J.A., Woods, G.L. & Aguilar, J.J. Location of equine oviductal embryos on day 5 post ovulation and oviductal transport time of day 5 embryos autotransferred to the contralateral oviduct. Theriogenology 46, 1477–1483 (1996).

11. Koester, H. Ovum transport, in Mammalian Reproduction 189–228 (Springer Berlin Heidelberg, 1970).

12. White, J.K. et al. Genome-wide generation and systematic phenotyping of knockout mice reveals new roles for many genes. Cell 154, 452–464 (2013).

13. Black, D.L. & Davis, J. A blocking mechanism in the cow oviduct. J Reprod Fertil 4, 21–26 (1962).

14. Gott, A.L., Gray, S.M., James, A.F. & Leese, H.J. The mechanism and control of rabbit oviduct fluid formation. Biol Reprod 39, 758–763 (1988).

15. Belve, A.R. & McDonald, M.F. Directional flow of fallopian tube secretion in the Romney ewe. J Reprod Fertil 15, 357–364 (1968).

16. Kanayama, K. & Osada, H. Relationship between changes in volume of the oviductal fluid in the ampulla and the descent of ovulated eggs from the ampulla to the isthmus in mice. J Int Med Res 28, 20–23 (2000).

17. Black, D.L., Duby, R.T. & Riesen, J. Apparatus for the Continuous Collection of Sheep Oviduct Fluid. J Reprod Fertil 6, 257–260 (1963).

18. Hino, T. & Yanagimachi, R. Active Peristaltic Movements and Fluid Production of the Mouse Oviduct: Their Roles in Fluid and Sperm Transport and Fertilization. Biol Reprod 101, 40–49 (2019).

19. Black, D.L. & Asdell, S.A. Transport through the rabbit oviduct. Am J Physiol 192, 63–68 (1958).

20. Edgar, D.G. & Asdell, S.A. The valve-like action of the uterotubal function of the ewe. J Endocrinol 21, 315–320 (1960).

21. Hamann, J. et al. International Union of Basic and Clinical Pharmacology. XCIV. Adhesion G protein-coupled receptors. Pharmacol Rev 67, 338–367 (2015).

22. Liebscher, I. et al. A tethered agonist within the ectodomain activates the adhesion G protein-coupled receptors GPR126 and GPR133. Cell Rep 9, 2018–2026 (2014).

23. Arac, D. et al. A novel evolutionarily conserved domain of cell-adhesion GPCRs mediates autoproteolysis. EMBO J 31, 1364–1378 (2012).

24. Coleman, J.L., Ngo, T. & Smith, N.J. The G protein-coupled receptor N-terminus and receptor signalling: N-tering a new era. Cell Signal 33, 1–9 (2017).

25. Wright, G.J. Signal initiation in biological systems: the properties and detection of transient extracellular protein interactions. Mol Biosyst 5, 1405–1412 (2009).

26. Martinez-Martin, N. et al. An Unbiased Screen for Human Cytomegalovirus Identifies Neuropilin-2 as a Central Viral Receptor. Cell 174, 1158–1171 e1119 (2018).

27. Bushell, K.M., Sollner, C., Schuster-Boeckler, B., Bateman, A. & Wright, G.J. Large-scale screening for novel low-affinity extracellular protein interactions. Genome Res 18, 622–630 (2008).

28. Yulia, A., Singh, N., Lei, K., Sooranna, S.R. & Johnson, M.R. Cyclic AMP Effectors Regulate Myometrial Oxytocin Receptor Expression. Endocrinology 157, 4411–4422 (2016).

29. Bohnekamp, J. & Schoneberg, T. Cell adhesion receptor GPR133 couples to Gs protein. J Biol Chem 286, 41912–41916 (2011).

30. Dickens, C.J. & Leese, H.J. The regulation of rabbit oviduct fluid formation. J Reprod Fertil 100, 577–581 (1994).

31. Tay, J.I. et al. Human tubal fluid: production, nutrient composition and response to adrenergic agents. Hum Reprod 12, 2451–2456 (1997).

32. Skarnes, W.C. et al. A conditional knockout resource for the genome-wide study of mouse gene function. Nature 474, 337–342 (2011).

33. Bianchi, E., Doe, B., Goulding, D. & Wright, G.J. Juno is the egg Izumo receptor and is essential for mammalian fertilization. Nature 508, 483–487 (2014).

34. Green, M., Bass, S. & Spear, B. A device for the simple and rapid transcervical transfer of mouse embryos eliminates the need for surgery and potential post-operative complications. Biotechniques 47, 919–924 (2009).

35. Bartholdson, S.J. et al. Semaphorin-7A is an erythrocyte receptor for P. falciparum merozoite-specific TRAP homolog, MTRAP. PLoS Pathog 8, e1003031 (2012).

36. Chong, Z.S., Ohnishi, S., Yusa, K. & Wright, G.J. Pooled extracellular receptor-ligand interaction screening using CRISPR activation. Genome Biol 19, 205 (2018).

