## Supplementary Information for "Control of fluid flow by Adgrd1 is essential for mammalian oviductal embryo transport"

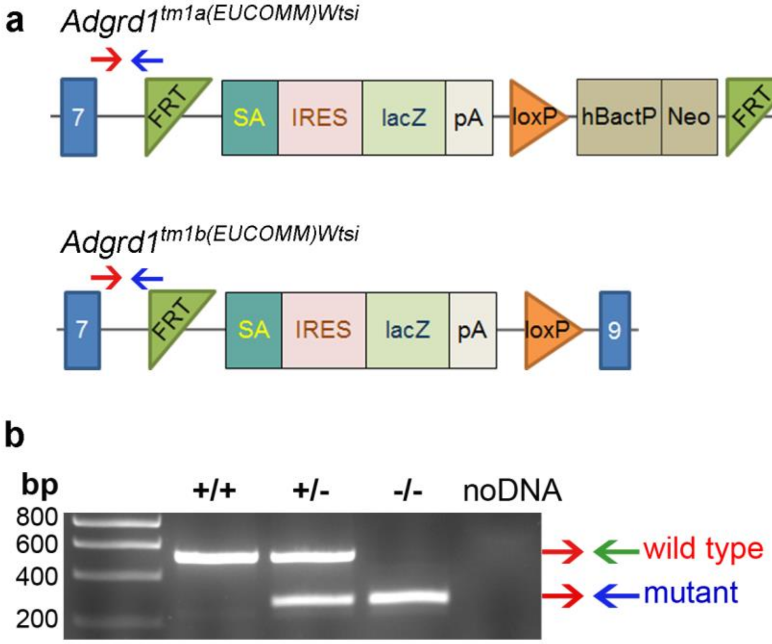


**Supplementary Fig. 1. Generation of *Adgrd1*$^{-/-}$ mice using a reporter-tagged deletion allele.** **a**, Targeted ES cells carrying the "knockout-first" tm1a allele were selected using standard techniques for homologous recombination. In the tm1a allele, *Adgrd1* function is disrupted by insertion of the LacZ marker gene and a floxed Neo selection cassette upstream of exon 8. Conversion to the LacZ-tagged null allele (tm1b) lacking exon 8 was obtained by crossing *Adgrd1*$^{tm1a(EUCOMM)WTSI}$ heterozygous females with *Hprt*$^{Tg(CMV\text{-}Cre)Brd}$ Cre recombinase-expressing males. The colony was maintained by mating heterozygous males and females. **b**, Mice were genotyped by PCR using the indicated primers to yield products of 512bp (wild-type allele) and 263bp (mutant allele).

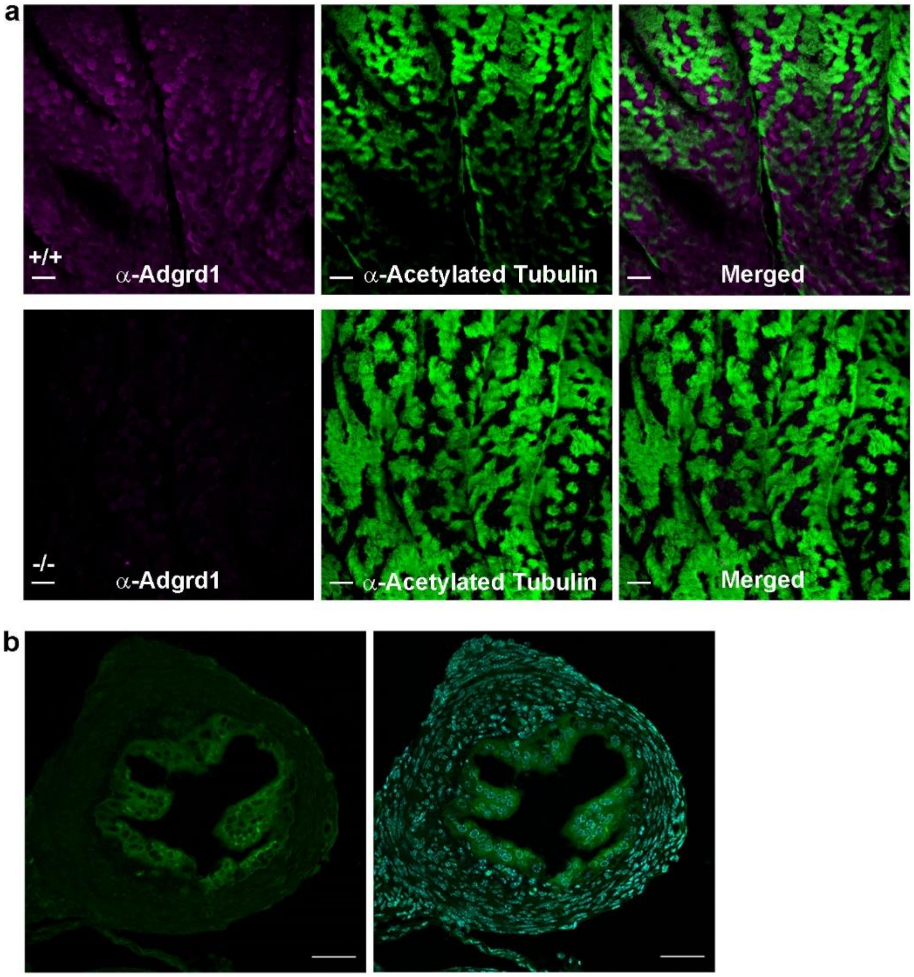


**Supplementary Fig. 2. Adgrd1 is expressed in ampullary and isthmic oviductal epithelial cells at different stages of the estrus cycle. a,** A rabbit polyclonal antibody raised against the entire extracellular region of mouse Adgrd1 specifically stains ampullary epithelium (purple) from wild-type (upper panels) but not *Adgrd*1-deficient mice (lower panels); anti-acetylated tubulin staining was used to mark ciliated cells (green). **b**, Expression of Adgrd1 in the epithelial cells of the isthmus is confirmed by immunostaining of oviductal sections with an antibody against the C-terminal fragment of human ADGRD1 (green, left panel) and nuclei counterstained with DAPI (blue, right panel). A representative example from a heterozygous female in diestrous is shown, scale bar represents 50 μm.

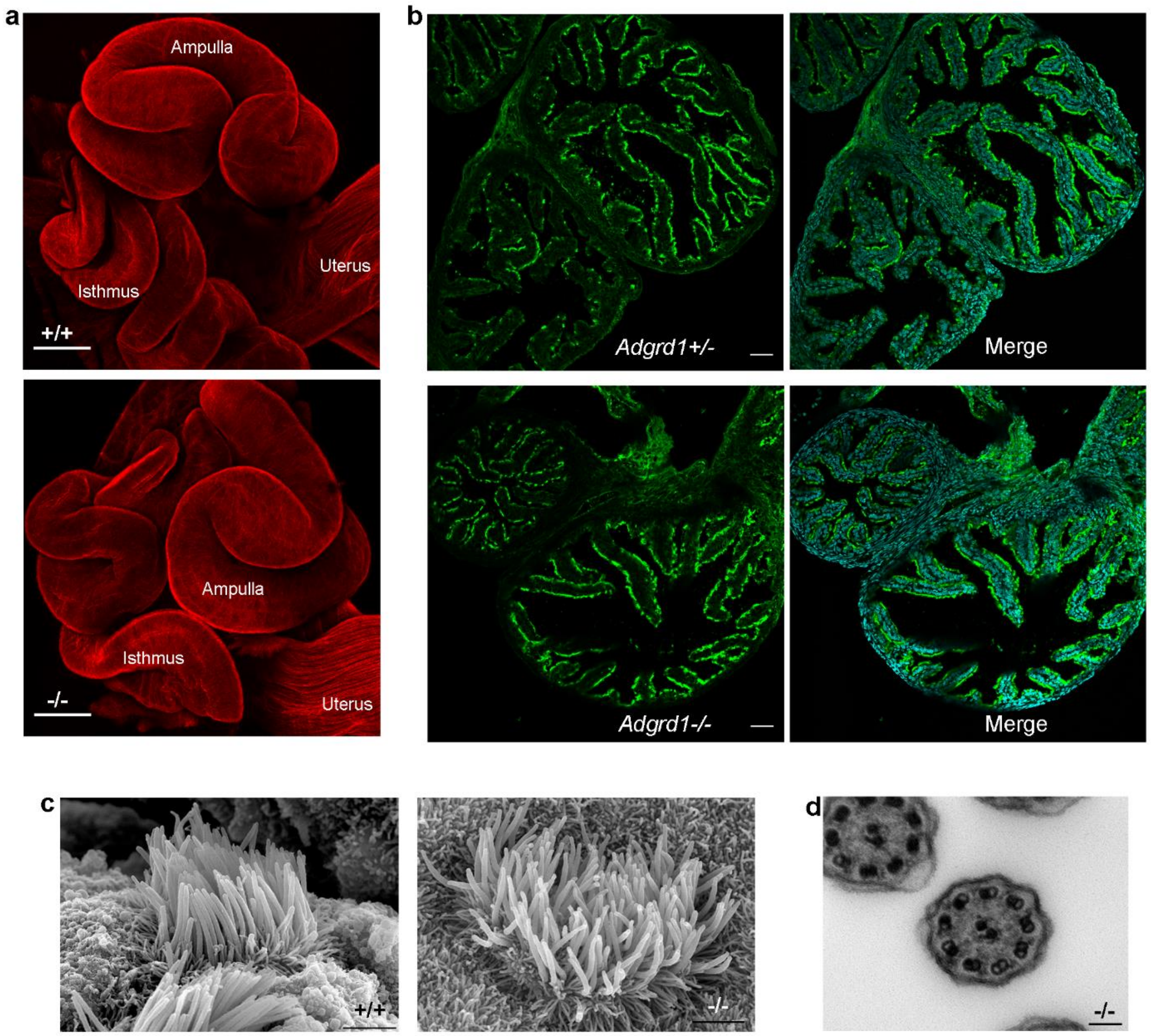


**Supplementary Fig. 3. Muscle and cilia appear normal in *Adgrd1*-deficient oviducts**. **a**, No overt defect is visible in the myosalpinx of *Adgrd1*$^{-/-}$ mice. Image shows 3D projection of oviducts from 17-day-old mice stained with a phalloidin-Texas Red conjugate, demonstrating no overt difference in muscle structure and organisation between wild-type (top panel) and mutant (lower panel); scale bar represents 250 µm. **b**, The distribution of ciliated cells is similar in the epithelium of control *Adgrd*$^{+/-}$ and *Adgrd1*$^{-/-}$ oviducts. Oviductal sections from adult females in diestrous were stained with an antibody against acetylated tubulin to mark cilia (green), and nuclei counterstained with DAPI (blue, right panel). Scale bar represents 50 µm **c**, Ciliated cells analysed by scanning electron microscopy did not differ in mutant *Adgrd1*$^{-/-}$ oviducts compared to wild-type controls; scale bar represents 1 µm. **d**,Transmission electron microscopy images of *Adgrd1*$^{-/-}$ cilia showed the usual 9+2 organisation of microtubules; scale bar represents 100 nm.

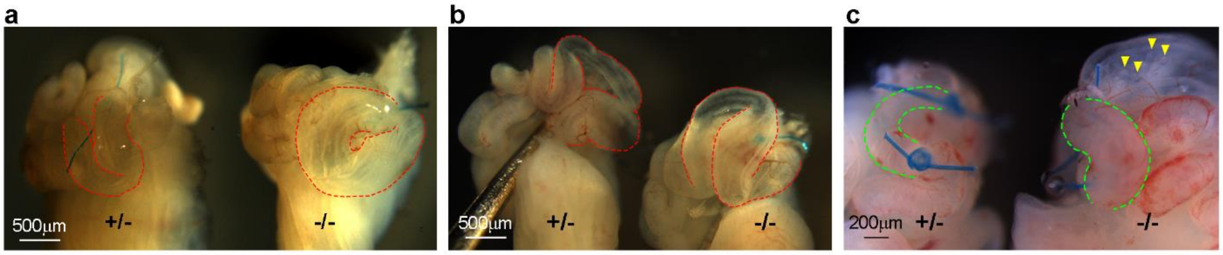


**Supplementary Fig. 4. *Adgrd1* regulates attenuation of post-ovulatory oviductal fluid production.** *Adgrd1* mutant (-/-) and heterozygous control (+/-) oviducts were ligated *in vivo* at E2.5 and collected four hours later; the ampullary region is highlighted by dotted red lines. **a**, and **b**, show representative examples of three independent experiments. Homozygous mutant oviducts show a more distended ampulla compared to the heterozygous control. **c**, Attenuation of oviductal fluid production is dysregulated in the isthmus of Adgrd1-mutant oviducts. Control heterozygous (left) and Adgrd1-deficient (right) oviducts at E1.5 were ligated in three different locations: the infundibulum, the ampullary-isthmic junction (AIJ), and within the isthmus just after the AIJ. The oviducts were collected four hours after ligation and images collected. The dotted green line shows a larger expansion of the isthmus in the mutant oviduct compared to control. The yellow arrowheads point to oocytes ectopically located in the ampulla of Adgrd1-mutant oviduct.

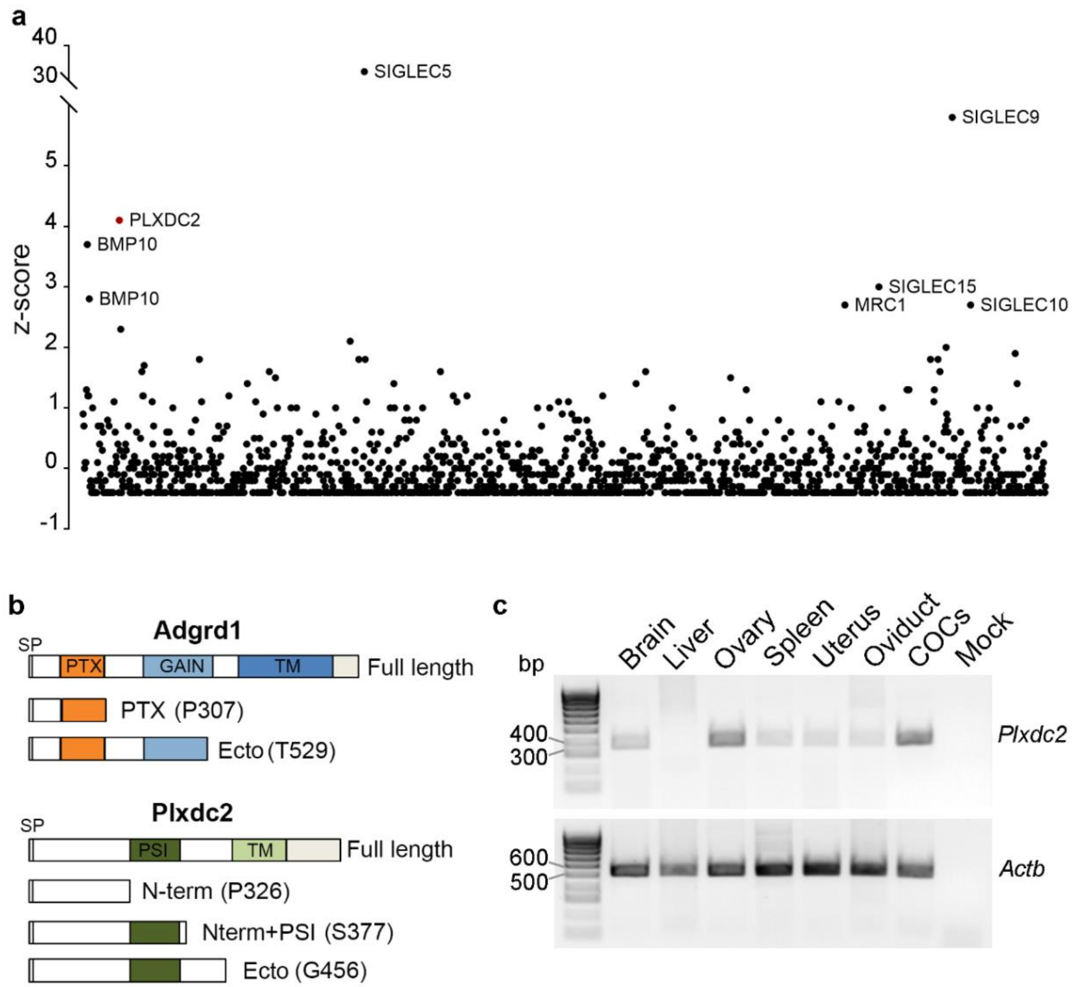


**Supplementary Fig. 5. A systematic high throughput ectodomain binding screen identified PLXDC2 as a ligand for ADGRD1. a**, A large panel of human receptor ectodomains was expressed as Fc-fusion proteins and immobilised on protein-A coated microtitre plates before testing for direct binding to a pentameric, β-lactamase-tagged ADGRD1 probe. Binding was quantified by measuring the absorbance of a hydrolysis product of β-lactamase at 485 nm. A z-score was calculated for each tested ligand and the identities of those ligands with a z-score > 2.5 are indicated. PLXDC2 was identified as a candidate ligand due to the high z-score and by comparing ligand behaviours with other binding probes in similar screens. Members of the SIGLEC (sialic acid-binding immunoglobulin-type lectins) family and MRC1 (Macrophage mannose receptor 1) are known glycan binding proteins which are often identified in these binding screens by directly interacting with common glycans present on the protein probe. The BMP10 ligands encode secreted proteins which were also repeatedly identified in previous screens irrespective of the protein binding probe presented and so not considered further. **b**, Schematic of mouse Adgrd1 and mouse Plxdc2 constructs. SP = signal peptide; PTX = pentraxin domain; GAIN = G-protein-coupled receptor (GPCR) autoproteolysis-inducing domain; TM = predicted transmembrane region; PSI = plexin-semaphorin-integrin domain. The amino acid identities and position of the exact truncation points are reported in parenthesis. **c**, RT-PCR showing the expression of *Plxdc2* in mouse tissues and cumulus-oocyte complexes (COCs).

**Supplementary Movie 1. Muscle function appears normal in *Adgrd1*-deficient oviducts.**

Beads and 2-cell embryos are regularly moved back and forth due to rhythmic muscular contractions within an *Adgrd1*$^{-/-}$ ampulla explant at E1.5. A contracting segment of the isthmus is visible in the upper left corner. The video is shown at four frames per second.

**Supplementary Movie 2. Absence of *Adgrd1* does not affect ciliary function on oviductal epithelium.**

The cilia that line the ampullary epithelium beat constantly in an *Adgrd1*$^{-/-}$ mutant oviduct. Effective transport is shown by movement of 15 µm microparticles along the ampullary folds.
